# CNN-based learning of single-cell transcriptomes reveals a blood-detectable multi-cancer signature of brain metastasis

**DOI:** 10.1101/2024.03.08.584083

**Authors:** Ryan Lusby, Debojyoti Chowdhury, Sarah Carl, Vijay K. Tiwari

## Abstract

Brain metastasis (BrM) is a serious complication of advanced cancers and remains difficult to predict before clinical symptoms appear. To investigate shared transcriptional features of BrM across tumour types, we integrated single-cell RNA sequencing (scRNA-seq) data from malignant epithelial cells derived from six carcinoma types, including lung, breast, colorectal, renal, ovarian, and melanoma. We applied ScaiVision, a supervised representation learning method, to classify tumour samples based on BrM status. The models achieved high predictive accuracy (area under the ROC curve > 0.90) across all six cancer types. This analysis identified a consistent multi-cancer gene expression signature associated with BrM, defined at single-cell resolution. To evaluate the clinical relevance of this signature, we assessed its presence in tumour-educated platelets (TEPs) from blood samples of patients with and without BrM. The signature was detectable in platelet RNA and distinguished patients with BrM from those without, indicating that features of the BrM-associated expression program are reflected in blood-derived material. These findings demonstrate that a transcriptional signature of brain metastasis can be identified across multiple tumour types using scRNA-seq and neural network-based analysis. The detectability of this signature in TEPs supports its relevance in a non-invasive context and provides a basis for further investigation into its utility for BrM risk assessment.

## Introduction

Brain metastases (BrM) are the most common intracranial tumours, occurring in approximately 20% of adult cancer patients^1,2^. They represent a devastating complication of systemic malignancy, with limited treatment options and poor survival outcomes^1,3^. Most BrMs arise from primary cancers of the lung, breast, melanoma, and other solid tumours, and their development often portends neurologic decline and dismal prognosis. Despite improvements in cancer care, effective means to predict, detect, or prevent BrM remain an unmet clinical need^3^. In particular, the absence of robust biomarkers for early identification of patients at risk for BrM hampers timely intervention and surveillance strategies. This clinical burden and diagnostic gap underscore the urgency for deeper biological insights into BrM pathogenesis and markers of metastatic potential.

Recent advances in single-cell RNA sequencing (scRNA-seq) have begun to characterise the cellular and molecular architecture of BrMs, although significant knowledge gaps persist^4,5^. Single-cell approaches allow high-resolution dissection of the tumour microenvironment, revealing diverse cell types and states that contribute to brain metastatic colonisation. Notably, emerging scRNA-seq studies have identified certain conserved stromal and immune features across BrMs from different cancers. For example, cancer-associated fibroblasts within brain metastatic lesions display a remarkably conserved activation profile characterised by abundant type I collagen and wound-healing signatures, mirroring fibrotic programs seen in primary tumours^6^. Similarly, metastasis-associated myeloid cells (including resident microglia and infiltrating macrophages) in BrM often converge on shared phenotypic states; for instance, coexisting pro-inflammatory (M1-like) and immunosuppressive (M2-like) macrophage subsets have been observed across brain metastases of various origins^7^. These conserved fibroblast and macrophage responses suggest that disparate tumour types co-opt parallel microenvironmental programs when colonising the brain. However, translating such biological insights into clinical diagnostics or therapies is not straightforward. In particular, the question remains how to leverage single-cell discoveries to identify predictive markers of BrM before metastases become clinically evident.

A multi-cancer approach to this question is especially valuable given that BrMs arise from multiple tumour lineages. To date, many studies of BrM have been siloed by cancer type, focusing, for example, on BrM in lung cancer or in breast cancer separately. While such studies have yielded important tumour-specific insights (for instance, single-cell analyses in lung adenocarcinoma have identified a distinct subpopulation of brain metastasis-associated tumour cells in the primary lung tumour, marked by genes such as S100A9, which correlates with subsequent BrM development^2^), they may overlook metastasis mechanisms that are shared across diseases. A comparative, cross-tumour analysis can reveal universal drivers and biomarkers of brain metastatic spread that single-tumour studies might miss. Indeed, evidence is mounting that certain BrM-associated gene expression changes in the tumour microenvironment (e.g. in stromal or immune compartments) are conserved irrespective of the cancer’s tissue of origin^6,7^. Identifying such common signatures could inform broadly applicable diagnostic tools. Recent pan-cancer single-cell atlases have characterised the cellular ecotypes and altered expression programmes of brain metastasis across tumour types^55^. Here we used a supervised model to derive a gene signature of BrM and tested whether it is detectable in blood.

In light of the above, the present study leverages scRNA-seq across multiple cancer types to identify BrM-associated markers in primary tumours. We used an interpretable deep learning framework, Scailyte AG’s in-house platform, ScaiVision to extract biologically meaningful features from scRNAseq data while maintaining model transparency. ScaiVision employs supervised representation learning to generate low-dimensional, yet biologically relevant, embeddings that distinguish BrM cells from their primary tumour counterparts. Through integrated gradients-based feature attribution analysis, our model identifies the individual genes that drive classification decisions, thus enabling direct biological interpretation of its output. This architecture allows for cross-tumour generalisation while preserving cell-type specific signals critical for the identification of metastasis-associated features. We further employed these genes to create a scoring system that uncovers cells responsible for brain tropism and analysed the plausible mechanism of action of these cells intrinsically as well as in coordination with the microenvironment. The resultant BrM signature was also detectable in liquid biopsies, specifically tumour-educated platelets, highlighting its potential as a non-invasive biomarker for early detection and risk stratification. More broadly, this strategy demonstrates how interpretable machine learning applied to single-cell data can uncover clinically actionable biomarkers and mechanistic insights into metastatic progression across cancer types, offering a blueprint for similar applications in other metastatic or treatment-resistant disease contexts.

## Results

### Modelling Brain Metastasis and Primary Cancers with Interpretable Neural Networks

Despite advancements in diagnostic techniques and therapeutic interventions, brain metastases (BrMs) remain associated with disproportionately high rates of morbidity and mortality. This clinical challenge underscores the urgent need to uncover robust molecular biomarkers that can facilitate early diagnosis, patient stratification, and development of targeted therapies across primary malignancies predisposed to cerebral dissemination. To address this unmet need, we designed a computational workflow centred on elucidating the transcriptional determinants of brain metastasis using single-cell transcriptomic profiles. Recognizing that conventional deep learning frameworks often function as “black boxes” and lack interpretability with respect to the biological relevance of features, we implemented a fully interpretable representation learning architecture tailored for single-cell classification. This framework enabled us not only to distinguish BrM samples from their primary tumour counterparts with high accuracy, but also to extract highly discriminative gene signatures that may define the metastatic phenotype at the single-cell level.

Our approach is grounded in the biological hypothesis that cells occupying transcriptionally similar states are likely to share functional and developmental programs, allowing us to infer the metastatic potential of individual cells based on gene expression similarities. Using this principle, we aimed to identify BrM-specific molecular programs and construct a brain metastasis signature enrichment score to evaluate their presence across individual cells (**Figure 1A**). We curated a dataset comprising single-cell RNA-sequencing (scRNA-seq) data from 115 patient samples, including 21 brain metastases and 94 primary tumours (**Figure 1B**, **Supplementary Figure 1A**). This dataset was divided into a training cohort (n=70; 13 BrM, 57 primary) and a validation cohort (n=45; 8 BrM, 37 primary) to support robust model development and independent performance assessment. To avoid bias from differences in cellular composition between primary and metastatic tumours, we restricted the model training to epithelial cells only (**Figure 1C**). Previous studies support the improved accuracy of epithelial cell-specific scRNA-seq signatures, emphasizing the importance of cell-type specificity in biomarker development^8^.

**Figure 1.**
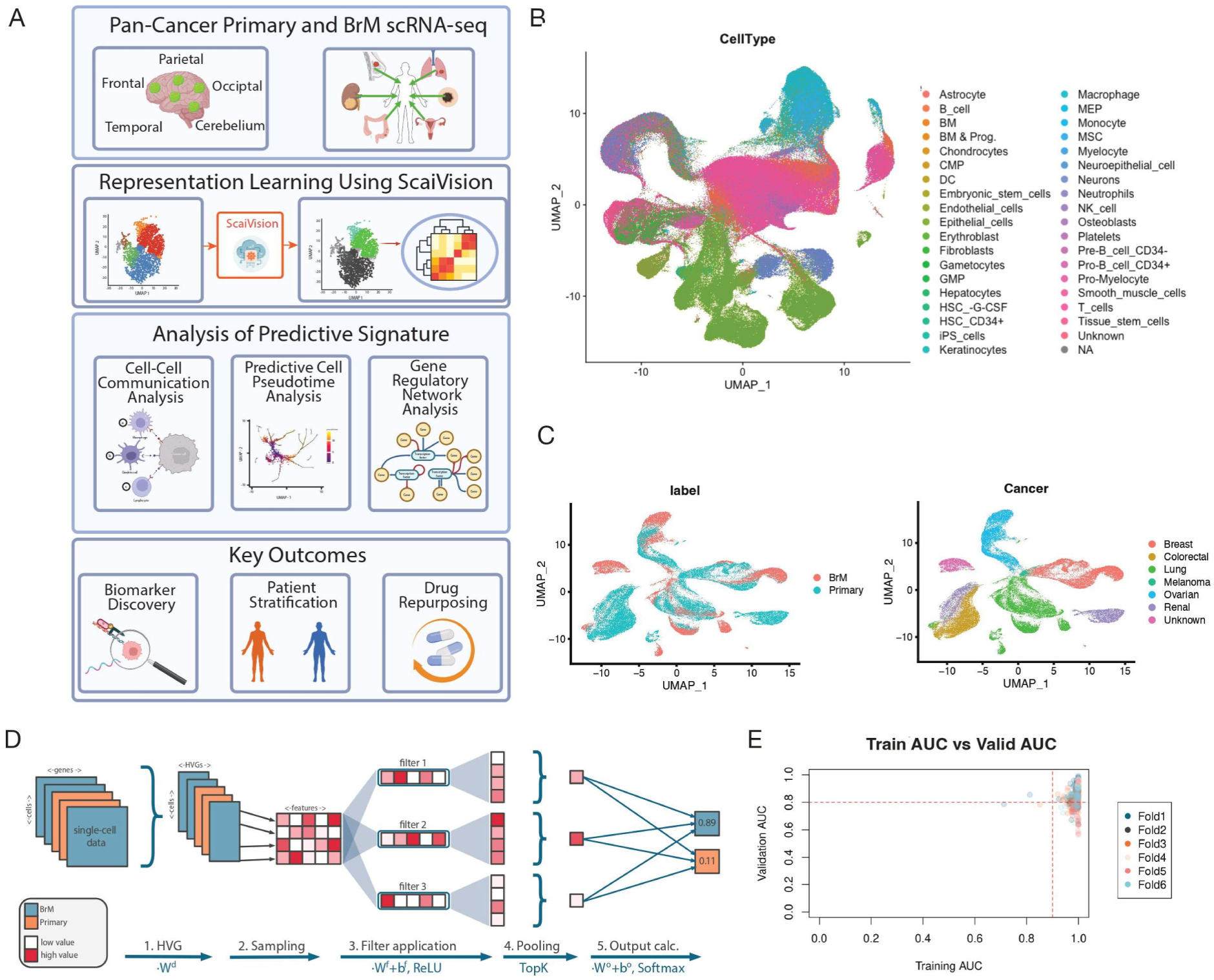
Identification and validation of a brain metastasis (BrM) signature in epithelial cells. **(A)** Workflow overview illustrating the strategy for identifying the BrM signature from single-cell RNA sequencing (scRNA-seq) data across multiple cancer types. **(B)** UMAP plot illustrating full multi-cancer dataset prior to subsetting and model training. Cells are coloured by cell type. Sample label and cancer type are shown in Supplementary Figure 1A. **(C)** Representative UMAP plots demonstrating epithelial cell selection for downstream analysis. Cells are coloured according to their cancer type origin from primary tumour or BrM samples. **(D)** Detailed schematic of the ScaiVision network architecture used for model training. **(E)** Scatter plots summarising Area Under the Curve (AUC) scores for the predictive models across training and validation datasets. The threshold (AUC > 0.9 and Validation AUC > 0.8) used for selecting high-performing models is indicated by a dashed line.

To initiate model training (**Figure 1D**), we first selected highly variable genes (HVGs) from the gene expression matrix, reducing dimensionality while retaining key variance associated with biological signal. From this representation, representative cells were randomly sampled to ensure diversity and avoid sampling bias. Subsequently, the data were processed through a series of convolutional filters, which scanned across features to learn localized transcriptional patterns indicative of BrM or primary origin. These filter activations were passed through a Rectified Linear Unit (ReLU) activation function to introduce non-linear transformations, followed by Top-K pooling, which retained the most informative features for classification optimizing model performance through extensive hyperparameter tuning^9^. The pooled features were then fed into a fully connected output layer with a SoftMax function, yielding class probabilities that distinguish BrM from primary samples. We trained a total of 300 deep learning models across six independent Monte Carlo cross-validation splits, each for up to 50 epochs, and retained the model weights corresponding to the epoch with the lowest validation loss to prevent overfitting. Evaluation metrics such as classification accuracy and cross-entropy log-loss demonstrated consistent performance and training stability across splits (**Supplementary Figures 1B-D**), attesting to the robustness and generalizability of our learning architecture.

For downstream biological interpretation, we selected models exhibiting high discriminatory performance, defined as Area Under the Receiver Operating Characteristic Curve (AUC) > 0.9 in the training set and AUC > 0.8 in the validation set (**Figure 1E**). Further evaluation was made including accuracy and cross-entropy log-loss. This stringent filtering yielded a final ensemble of 223 predictive models (**Figure 1C**; **Supplementary Figure 1B-D**), each capable of reliably identifying transcriptional features associated with metastatic potential. The high predictive accuracy of these models, coupled with their interpretability, lays a strong foundation for dissecting the molecular programs underpinning BrM.

### Identification of Gene Signature with High Accuracy in Prediction of BrM from Epithelial cells

To uncover the molecular drivers underpinning model predictions, we performed a systematic feature attribution analysis using Integrated Gradients (IG), implemented via the Captum.ai interpretability library^10^. This approach enables a principled estimation of each input gene’s contribution to the model’s output by integrating gradients of the prediction probability with respect to the input features. We applied IG across our ensemble of high-performing deep learning models to generate an aggregated attribution matrix that prioritizes genes based on their relevance in distinguishing brain metastasis (BrM) from primary tumour cells. Through this aggregation, we derived a consensus ranking of genes based on their mean attribution scores, ultimately identifying 173 highly variable genes (HVGs) with consistently high explanatory power across models. The top 20 most influential genes from this list are visualized in **Figure 2A**, while the complete ranked gene set is provided in Supplementary Table 2. We collectively refer to these genes as the BrM signature, representing a core set of transcriptional features that may be mechanistically or functionally linked to brain metastatic progression and therefore warrant further experimental validation and therapeutic exploration.

**Figure 2.**
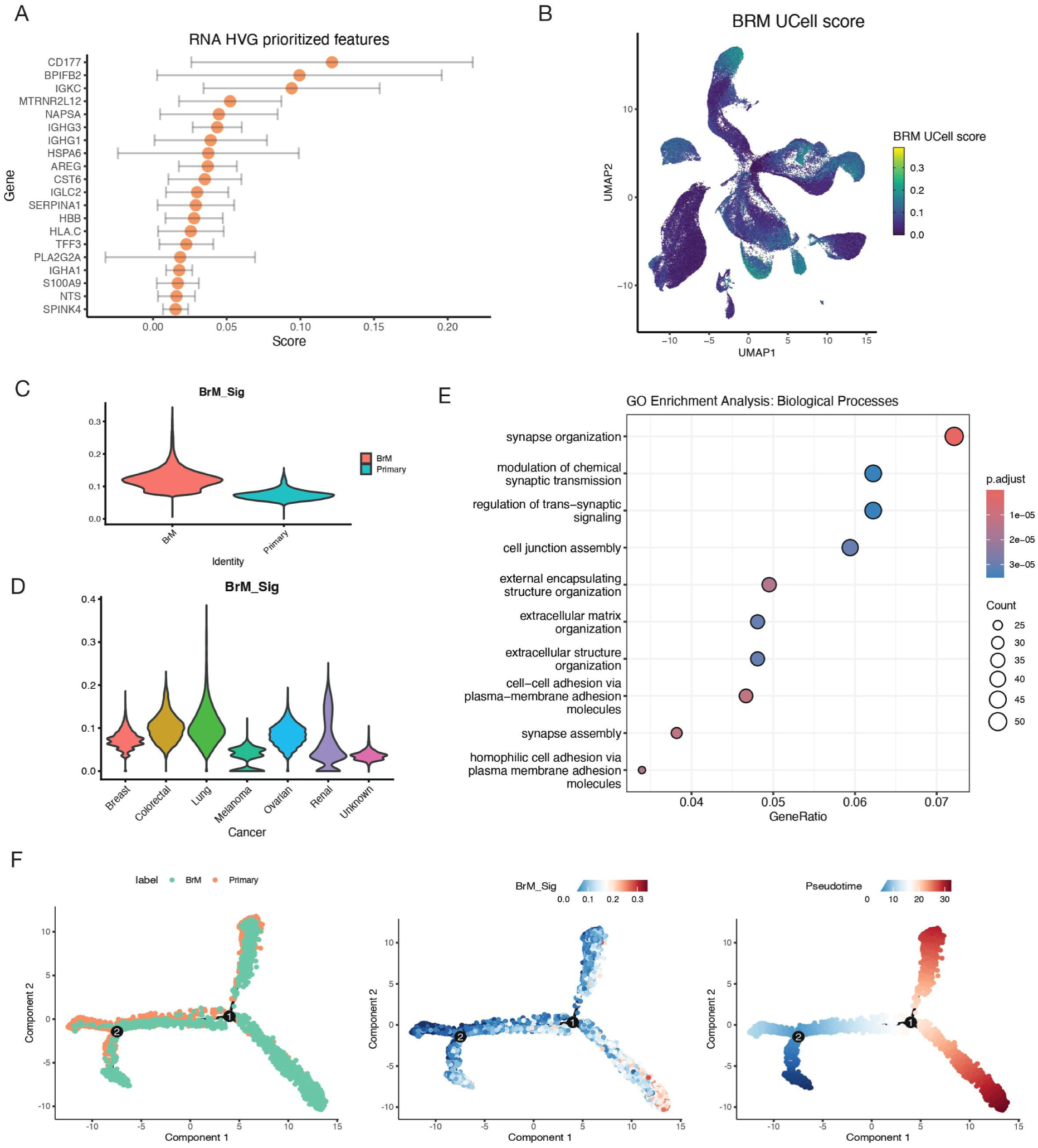
Prediction of differential features of BrM using Machine Learning. **(A)** Chart showing the top 20 genes ranked by Integrated Gradients attribution scores, highlighting their contribution to model predictions. These genes represent the core components of the identified BrM signature (complete list provided in Supplementary Table 2). Points show the mean scores and error bars indicate the standard deviation of scores across models (n =223). **(B)** UMAP projection of epithelial cells, coloured according to BrM signature scores calculated using UCell. Higher scores indicate greater inferred metastatic potential, highlighting cell populations from confirmed BrM samples and subsets within primary tumours. **(C)** Violin plots comparing BrM signature scores (UCell) for epithelial cells between primary and BrM samples. Each violin represents all epithelial cells from the corresponding sample type. **(D)** Violin plots comparing BrM signature scores (UCell) for epithelial cells among cancer types. **(E)** Gene Ontology (GO) enrichment analysis illustrating biological processes significantly enriched in high-scoring epithelial cells (top 20%) compared to low-scoring epithelial cells (bottom 20%), derived from differential pseudobulk gene expression analysis. Dot size indicates gene count, and colour gradient represents adjusted p-values. Notably enriched pathways include extracellular matrix organisation, cell-cell adhesion, and synapse-related processes linked to metastatic capability. **(F)** Monocle 2 trajectory plot showing inferred pseudotime progression for cells from paired primary lung and BrM samples. Cells are coloured by sample origin, BrM signature score (UCell) and inferred pseudotime.

To assess the biological relevance and predictive utility of the BrM signature across individual cells, we computed a relative metastatic potential score for each epithelial cell, spanning both BrM and primary tumour origins, using the UCell scoring framework, a rank-based enrichment scoring method optimized for single-cell datasets^11^ (**Figure 2B**). As anticipated, epithelial cells derived from histologically confirmed brain metastases exhibited consistently high metastatic potential scores, validating the sensitivity of our signature to known metastatic phenotypes (**Figure 2C**). Importantly, a subset of epithelial cells within primary tumour samples also received high metastatic potential scores, suggesting the presence of transcriptionally primed subpopulations with latent metastatic capabilities (**Figures 1C, 2B and 2C**). This finding reinforces the notion that metastatic potential can be heterogeneous within primary tumours and highlights the potential of our signature to prospectively identify high-risk cell populations prior to dissemination. Moreover, the BrM signature demonstrated robust predictive capacity across multiple cancer types (**Figure 2D**), further supporting its cross-tumour applicability and biological generalizability. Within each cancer type, BrM samples exhibited higher median scores than primary tumour samples (Supplementary Figure 1E), reaching statistical significance in breast, melanoma and ovarian cancer (Wilcoxon rank-sum test with Benjamini-Hochberg correction; adjusted p = 0.001, 0.016 and 0.032, respectively). Comparisons in colorectal and renal cancer (one BrM sample each) and in lung cancer (two BrM samples) were underpowered, and the difference in lung cancer was small (median z-scores of 1.76 and 1.41 for BrM and primary samples, respectively). To further stratify by metastatic potential, cells were binned into high-scoring (top 20%), mid-scoring (20-80%), and low-scoring (bottom 20%) groups. Differential pseudobulk gene expression analysis comparing high- and low-scoring cells identified significant transcriptional differences associated with elevated metastatic potential (**Figure 2E**), including upregulation of pathways related to extracellular matrix remodelling, extracellular structural organization, synaptic signalling, and cell adhesion in cells with high metastatic scores (**Figure 2E**). These pathways are well-known contributors to metastatic processes, underscoring the biological validity of our signature.

To capture the dynamic progression from primary to metastatic states, we analysed paired scRNA-seq data from primary lung tumours and matched brain metastases from two patients^12^, enabling direct assessment of our BrM signature across the metastatic continuum. These paired samples were not included in the model training or validation datasets. Following standard preprocessing, BrM scores were computed for each cell using our established gene signature. As expected, scores were elevated in brain metastases, particularly within epithelial cells (**Supplementary Figure 2A-B**). To map the trajectory of metastatic progression, we employed pseudotime analysis using Monocle2^13^. Cells were embedded using DDRTree, revealing a pseudotemporal axis that strongly correlated with BrM scores, further validating the signature’s ability to capture transcriptional progression from primary to metastatic states (**Figure 2F**). To pinpoint genes with dynamic regulation across this trajectory, we applied GeneSwitches R package^14^, identifying key genes activated or repressed over pseudotime. Genes such as ZBTB16, NFIA, NKD1, and SOX6 were switched on, while DOCK2, IGHA1, and MUC1 were switched off (**Supplementary Figure 2C**). These genes include transcription factors and surface proteins with potential roles in metastatic transition and niche adaptation.

Gene ontology analysis of these temporally regulated genes revealed enrichment of biological processes including cell proliferation, adhesion, lymphocyte activation, and immune development. Notably, immune-related pathways, such as B-cell receptor signalling and phagocytic engulfment, were selectively upregulated during later stages of pseudotime, implicating them in the final stages of metastatic establishment within the brain (**Supplementary Figure 2D**).

### VEGF Signalling Correlates with BrM Score and Distinguishes Cell-Cell Networks in Primary Tumours and Brain Metastases

To broaden the scope of our analysis, we applied the BrM signature across the entire single-cell dataset, which includes a range of tumour and stromal cell types. This confirmed high BrM scores in cells from established brain metastases but also identified high-scoring cells within primary tumours, supporting the signature’s potential to flag cells at risk of metastasis even before dissemination. High-scoring cells were found across multiple cancer types (**Supplementary Figures 3A-C**), and across diverse cell populations including monocytes, endothelial cells, neuroepithelial cells, and neutrophils. Following our established stratification (top 20%, middle 60%, bottom 20%), differential pseudobulk gene expression analysis revealed distinct transcriptional programs in high-scoring cells. These were enriched in pathways related to extracellular matrix remodelling, immune modulation, chemotaxis, and cell adhesion (**Supplementary Figure 3D**), processes closely tied to metastatic progression. As the signature was derived from epithelial cells, this analysis across the whole dataset should be considered exploratory, and the comparison of high- and low-scoring cells was not adjusted for cell type composition.

Recognizing the BrM signature’s ability to identify metastatic potential across both tumour and stromal compartments, we next investigated cell-cell communication networks that might drive this phenotype. Tumour-stroma communication is well-documented as a key factor in cancer metastasis and treatment response, particularly within the complex microenvironment of the brain^15–17^. Using CellChat^18^ to identify the differential number of interactions between high- vs low-scoring cells, we found that several signalling pathways were significantly upregulated in high-scoring cells, with ANGPT emerging as the most enriched. VEGF signalling was also markedly more active in high-scoring cells compared to low-scoring ones (**Figure 3A**). ANGPT, CDH1, CADM and IGF signalling was detected exclusively in high-scoring cells. VEGF was selected for further investigation on the basis of its established role in brain metastasis and its support by several independent analyses described below (ligand-receptor inference with CellChat and LIANA, gene regulatory network analysis, spatial transcriptomics and drug repurposing); the remaining pathways were not investigated further in this study. Endothelial cells were key players in this communication, particularly through the VEGFA-VEGFR1 ligand-receptor axis, known to be associated with metastatic progression in various cancers^19–21^ (**Supplementary Figure 3E**). To validate these findings, we used LIANA (LIgand-receptor ANAlysis tool^22^), a consensus-based tool aggregating multiple cell-cell interaction algorithms.

**Figure 3.**
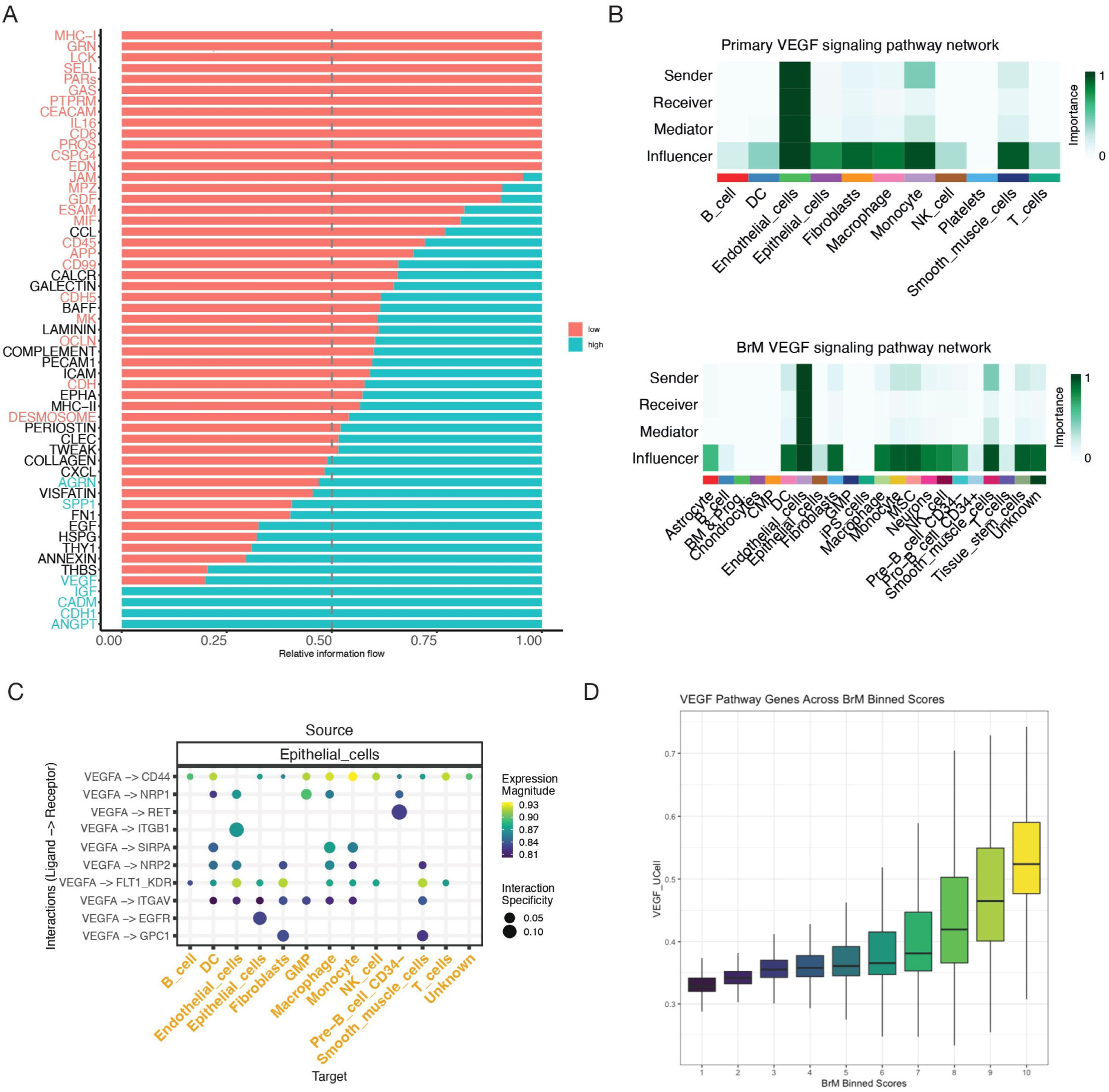
BrM signature scoring and cell-cell communication network analysis across the multi-cancer dataset. **(A)** CellChat pathway analysis comparing signalling pathway activity between high-scoring and low-scoring cells. Bar size represents relative pathway strength between the two conditions; colour indicates activity in high or low BrM scored cells (red=high, blue=low). Significant pathways are coloured on the y-axis **(B)** CellChat network heatmaps comparing the architecture of VEGF signalling pathways specifically within high-scoring cells derived from primary tumour sites and BrM sites. **(C)** Plot showing significantly enhanced activity scores for VEGF ligand-receptor pairs, particularly VEGFA-VEGFR1, in high-scoring cells compared to low-scoring cells, based on LIANA analysis. **(D)** Box plot showing UCell scores for a literature-derived VEGF signalling target gene set across cells binned into deciles based on their BrM signature score.

LIANA analysis confirmed the dominant role of epithelial cells as senders and endothelial cells as receivers in high-scoring populations and again highlighted VEGFA-VEGFR1 as a central interaction (**Figures 3B-C**).

To determine whether communication patterns varied by tumour site, we separately analysed high-scoring cells from primary tumours and BrM samples. While VEGF signalling remained a consistent feature across both contexts, the structure of the interaction networks shifted. In primary tumours, epithelial cells played a more prominent signalling role, whereas dendritic cells were more influential within BrM sites (**Figure 3B**). These findings point to a dynamic reconfiguration of VEGF-related communication networks as metastases establish in the brain.

Finally, we explored the transcriptional dynamics of VEGF target genes during metastatic progression. Using literature-derived VEGF targets^23–25^ and UCell scoring, we observed a gradual increase in VEGF target expression across cells binned by BrM score (in 10% increments), with expression peaking in the highest-scoring group (**Figure 3D**). This trend was supported by a strong correlation between VEGF signalling scores and BrM scores (R = 0.7, p < 2.2e-16) (**Supplementary Figure 3F**). These results underscore the importance of VEGF signalling not just as a marker, but potentially as a functional driver of metastatic transition. Together, our findings suggest that VEGF-mediated tumour-stroma interactions, particularly via VEGFA-VEGFR1, are critical in supporting the emergence and survival of metastatic cells in the brain.

### Gene Regulatory Network Analysis Identifies ETS1 as a Putative Regulator of VEGF-driven Brain Metastasis

VEGF signalling has been consistently implicated in angiogenesis and metastatic progression, and our previous analyses suggest its involvement across diverse tumour lineages in brain metastasis. To investigate the regulatory control underpinning this VEGF-driven BrM phenotype, we examined transcriptional regulatory networks that may govern VEGF pathway activity during metastatic progression.

We analysed single-cell RNA-seq data from two patients with matched primary lung tumours and brain metastases, previously used in our pseudotime analyses. Cells were stratified into nine equal-sized bins according to their BrM signature score distribution, allowing us to trace regulatory changes across a continuum of BrM potential. Gene regulatory networks (GRNs) were independently inferred for each bin using CellOracle^26^, enabling the identification of transcription factors (TFs) with differential connectivity as a function of metastatic progression. In the highest BrM score group, where VEGF pathway gene expression was also most pronounced, TFs such as MYC, STAT1, MEF2C, and notably ETS1 demonstrated high degree centrality (**Figure 4A**), indicating their potential importance in coordinating pro-metastatic transcriptional programs. Among these, ETS1 emerged as a particularly compelling candidate due to its well-established role in regulating VEGF-mediated angiogenesis and tissue invasion. By comparing TF connectivity across bins, we observed a progressive increase in ETS1 regulatory influence, paralleling VEGF pathway activation and BrM score enrichment (**Figure 4B**).

**Figure 4.**
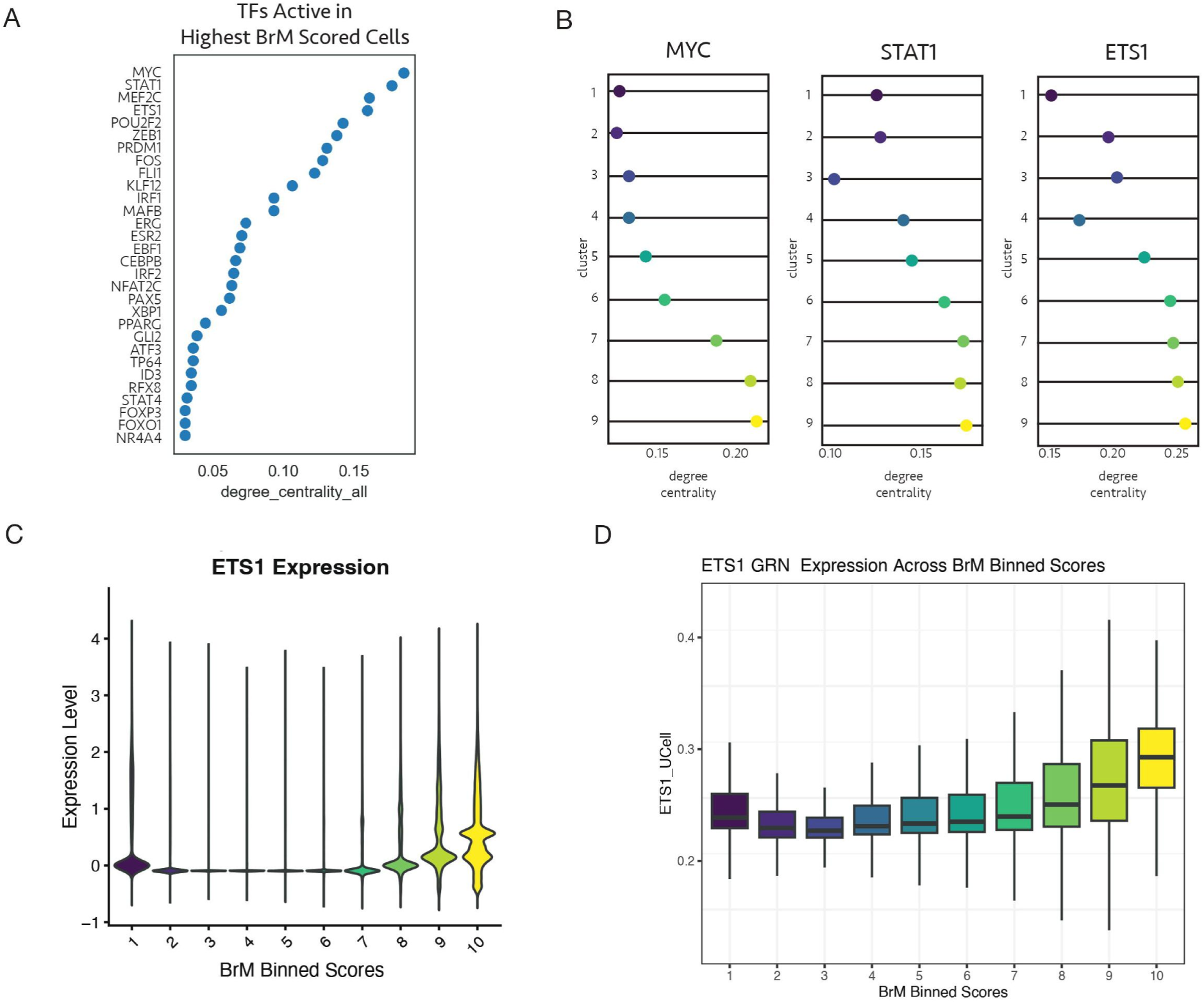
Gene regulatory network analysis implicates VEGF signalling drivers. **(A)** Dot plot illustrating transcription factor (TF) degree centrality within the gene regulatory network (GRN) inferred by CellOracle for the highest BrM score bin from paired primary/BrM samples. **(B)** Line plots showing the change in network connectivity (degree centrality) for key TFs (MYC, STAT1, ETS1) across nine BrM score bins. **(C)** Violin plot showing ETS1 gene expression distribution across BrM score bins in the multi-cancer scRNA-seq dataset. **(D)** Box plots showing UCell scores for ETS1 target genes (derived from CellOracle GRN) across BrM score bins in the multi-cancer scRNA-seq dataset.

We next focused on characterising ETS1 activity in the broader multi-cancer dataset to assess its relevance in a VEGF-driven BrM context. Although ETS1 transcript abundance was modest, consistent with known limitations of scRNA-seq in capturing TFs, its expression levels increased with higher BrM scores (**Figure 4C**). To better quantify its regulatory activity, we extracted ETS1 target genes from the inferred CellOracle networks and calculated a UCell enrichment score for each cell based on the expression of these downstream targets. This GRN-derived ETS1 activity score exhibited a clear upward trajectory across BrM score bins (**Figure 4D**), suggesting that ETS1 not only correlates with VEGF-driven metastatic potential, but may also act as a functional regulator of the transcriptional programs underlying this phenotype.

Together, these findings highlight ETS1 as a putative transcriptional regulator of VEGF-associated gene networks in brain metastasis. By integrating regulatory network inference with activity scoring and expression profiling, we reveal a layered regulatory architecture that underlies the VEGF-driven BrM phenotype. This approach allows us to sensitively detect key transcriptional drivers, even when expression alone is insufficient, and underscores the value of network-based strategies for dissecting the regulatory basis of metastasis to the brain.

### Spatial Transcriptomics Reveal VEGF-Driven Microenvironments in BrM Regions

To expand our transcriptomic insights, we analysed spatial transcriptomics data derived from four patients diagnosed with colorectal brain metastases. This approach enabled us not only to validate the BrM signature directly within the tissue architecture^27^ but also to investigate how metastatic potential varies across distinct microanatomical regions of the tumour and improve our understanding of metastasis biology^28^.

We applied UCell scoring to map BrM signature activity across the entirety of each spatial transcriptomic slide. In the patient shown, BrM signature scores were highest in inflammatory, vessel-containing and tumour-adjacent regions and lowest within the tumour region itself, indicating that high scores were not confined to the tumour core. The spatial pattern differed among the other three patients: scores were similar across most regions in one, elevated in blood-associated regions in another, and highest in a tumour region in the fourth (**Figure 5A**, **Supplementary Figure 4A**). This spatial distribution revealed pronounced heterogeneity in metastatic potential, underscoring the complex microenvironmental landscape that shapes tumour progression. The high scores in inflammatory and vessel-associated regions are consistent with those observed in endothelial and myeloid cells in the single-cell data. However, because each Visium spot contains multiple cells, spot-level scores cannot be attributed to a specific cell type.

**Figure 5.**
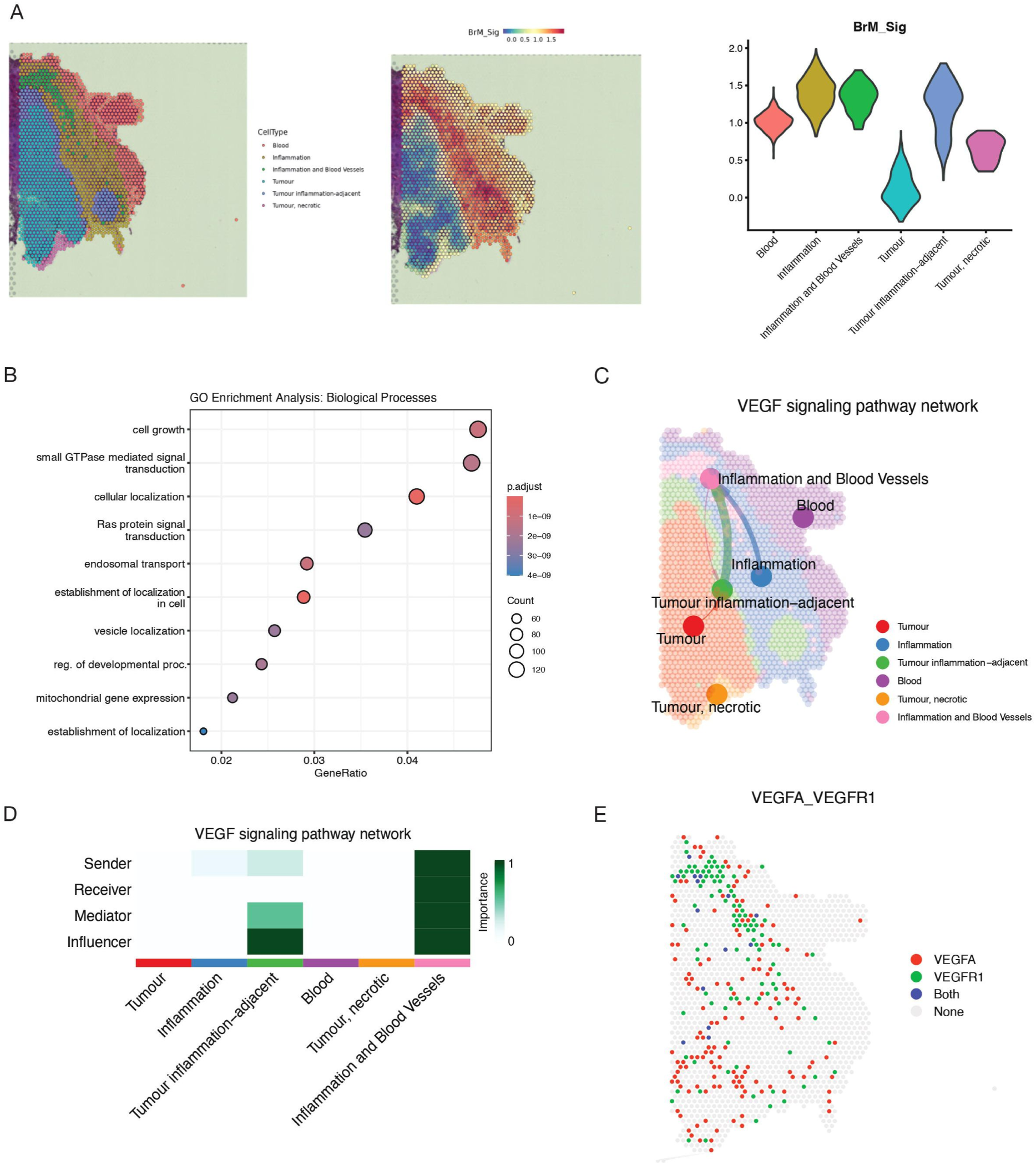
Spatial transcriptomics analysis reveals spatial organization of BrM potential and VEGF signalling. **(A)** Spatial feature plots showing BrM signature scores (UCell) mapped onto tissue section of a colorectal cancer BrM patient. Colour scale indicates BrM score. Key anatomical regions are annotated. Violin plots show BrM signature scores for each annotated region. **(B)** Gene Ontology (GO) enrichment analysis results for biological processes upregulated in high-scoring regions (top 20% BrM score) compared to other regions, based on spatially-resolved differential gene expression analysis. Dot size represents gene count; colour indicates significance level. **(C)** CellChat VEGF network plots projecting inferred VEGF pathway activity scores onto tissue sections. Arrow direction indicates signalling flow; line width indicates interaction strength. **(D)** CellChat network heatmap plot summarizing inferred VEGF signalling interactions between annotated tissue regions. **(E)** Spatial feature plots showing VEGFA and VEGFR1 expression mapped onto tissue sections. Spots are coloured according to expression of VEGFA, VEGFR1, both or neither.

To further dissect molecular differences underpinning these variations, we performed differential gene expression analysis comparing high-scoring regions (top 20% BrM signature activity) to all other tumour areas. This analysis identified enrichment of pathways involved in mitochondrial function, endosomal transport, vesicle localization, and Ras signalling (**Figure 5B**). These findings highlight active cellular processes potentially driving or sustaining metastatic behaviour within these spatially distinct niches.

Given the well-established role of VEGF signalling in promoting brain metastasis, we next utilised CellChat to characterize spatial patterns of intercellular communication. VEGF-mediated interactions were consistently prominent across all patient samples, with signalling concentrated in inflammation- and blood vessel-associated regions adjacent to the tumour in the patient shown, whereas tumour regions were among the principal sources of signalling in the other three patients (**Figure 5C-D**, **Supplementary Figure 4B**). Notably, in the patient shown, these VEGF-enriched zones strongly overlapped with regions exhibiting elevated BrM signature scores (**Figure 5A**), reinforcing the link between VEGF signalling and enhanced metastatic capacity. To visualize these dynamics in situ, we projected VEGF pathway activity directly onto spatial maps of tumour sections. This projection revealed pronounced enrichment of VEGF signalling in tumour-adjacent vascular niches and inflammation-associated areas, microenvironments critical for tumour-stromal crosstalk and likely contributors to metastatic progression (**Figure 5D**, **Supplementary Figure 4C**).

Additionally, we examined the spatial distribution of the key VEGF signalling ligand-receptor pair VEGFA-VEGFR1, which we previously identified in the pan-cancer scRNA-seq data. Spatial mapping showed that VEGFA and VEGFR1 were expressed predominantly in distinct spots, with few spots co-expressing both, as expected for a secreted ligand acting on neighbouring receptor-expressing cells (**Figure 5E**). This suggests that VEGFA-VEGFR1 signalling plays a pivotal role in supporting tumour cell survival, angiogenesis, and dissemination within the brain metastatic niche, as a central axis that facilitates metastatic progression^29^.

### Drug repurposing analysis identifies Pazopanib as a potential therapeutic candidate targeting VEGF-driven brain metastasis

To identify potential therapeutic compounds capable of reversing the transcriptional programs associated with high metastatic risk, specifically those linked to VEGF-driven brain metastasis, we employed the ASGARD R package^30^ on our integrated multi-cancer single-cell RNA-seq dataset. Cells were stratified according to their BrM scores, with the top 20% defined as high-BrM and the bottom 20% as low-BrM. Differential gene expression analysis was then conducted using limma, comparing high-versus low-scoring cells to identify genes consistently up- or downregulated in the metastatic state.

The resulting differentially expressed genes served as input to ASGARD’s drug repurposing pipeline, which mines the extensive L1000 drug-response dataset, comprising over 590,000 compound profiles, to identify small molecules capable of reversing the metastatic gene expression signature. Compounds were ranked based on their predicted efficacy in modulating BrM-associated gene expression, with candidates filtered by a false discovery rate (FDR) threshold of <0.05. Promising drug candidates with a predicted drug score >0.5 and FDR <0.05 were identified across a range of malignancies, including breast, colorectal, lung, ovarian, and renal (**Figure 6A**). The screen was not restricted to VEGF-related genes, and the predicted candidates also included compounds acting through other pathways, such as crizotinib, vorinostat and tretinoin (Figure 6A). Pazopanib was selected for further analysis because it targets the VEGF axis implicated in the preceding analyses; the remaining candidates were not investigated further in this study.

**Figure 6.**
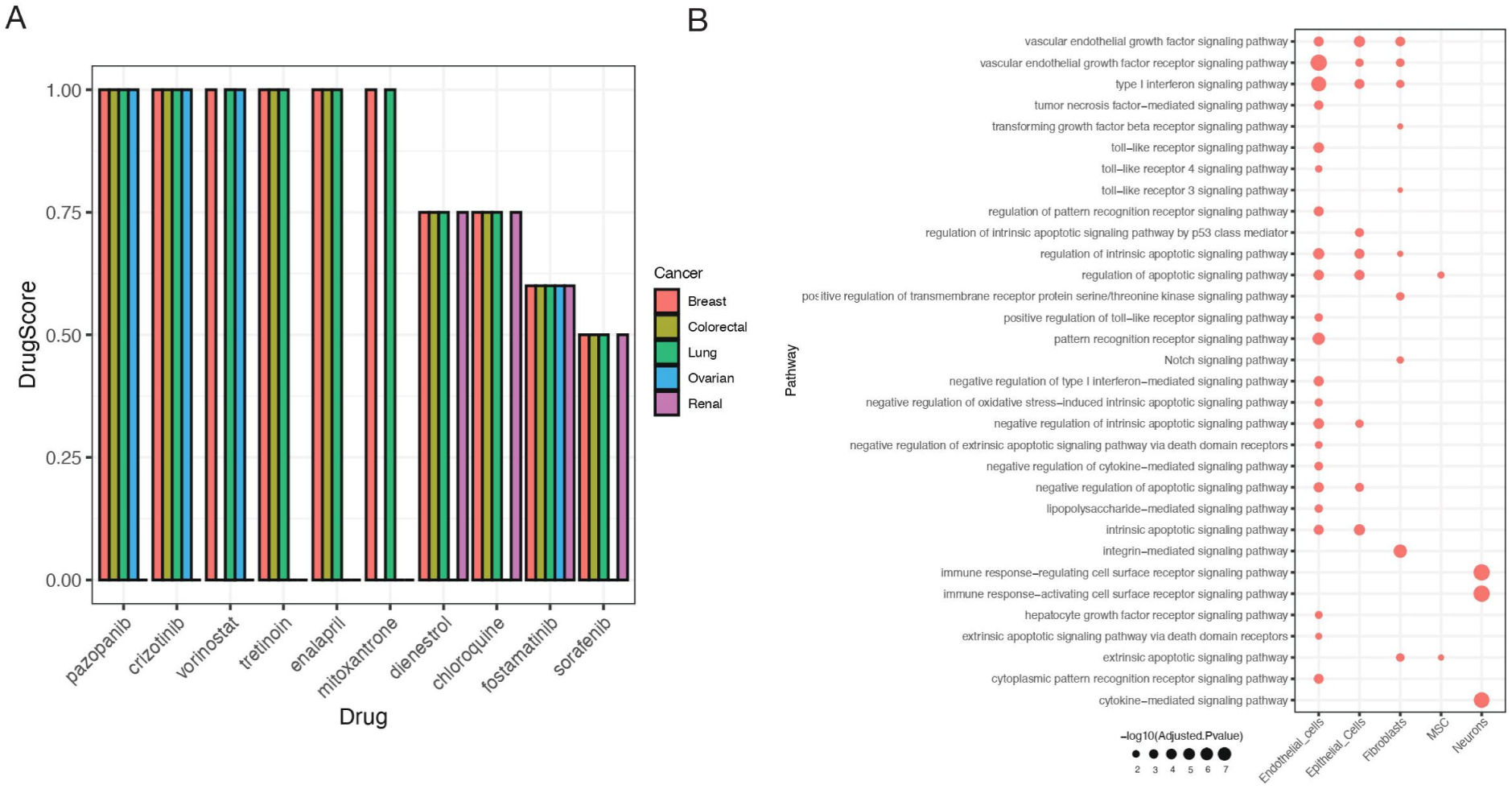
Pazopanib as a repurposing candidate for targeting VEGF signalling in brain metastasis. **(A)** Bar plots summarizing ASGARD drug repurposing results. Y-axis indicates the predicted drug score (reversal potential). **(B)** Gene ontology enrichment of predicted Pazopanib target genes (identified by ASGARD) within high BrM-scoring cells across different cell types.

Among the top-ranking candidates was Pazopanib, a multi-kinase inhibitor with known clinical activity in metastatic cancer, and a well-established inhibitor of VEGF receptors^31^. Given our earlier findings implicating VEGF signalling as a key driver of the BrM phenotype, we further examined Pazopanib’s mechanism of action across cancer types in our dataset. Using ASGARD, we extracted the predicted Pazopanib-responsive genes in high BrM-scoring cells and mapped these targets onto cell types and signalling pathways (**Figure 6B**). Pazopanib’s regulatory footprint included modulation of VEGF-associated gene expression in epithelial tumour cells, endothelial compartments, and stromal fibroblasts, indicating potential to disrupt both tumour-intrinsic and microenvironmental drivers of metastasis. Further experimental validation is required to assess its efficacy in BrM-specific models, and to explore potential synergy with existing therapies. By directly targeting the transcriptional underpinnings of VEGF-associated metastatic programs, Pazopanib could serve as a precision therapeutic agent to mitigate the risk or progression of brain metastasis across multiple tumour types.

### BrM Scoring Is Detectable in Tumour-Educated Platelets of BrM patients

Given the clinical need for non-invasive, real-time biomarkers to monitor metastatic disease progression, we sought to determine whether our brain metastasis (BrM) gene signature could be detected in peripheral blood samples from cancer patients. Liquid biopsy approaches, such as blood-based transcriptomic profiling, offer several advantages over tissue biopsies, including minimal patient discomfort, feasibility for repeated sampling, and the potential for dynamic disease monitoring over time^32,33^.

To investigate this, we leveraged a publicly available bulk RNA-sequencing dataset derived from tumour-educated platelets (TEPs) collected from patients representing six distinct cancer types. This cohort included individuals with and without metastatic disease, allowing us to compare transcriptional profiles between these two clinical states. As ScaiVision is designed for single-cell input, and training a model on bulk data would require considerably more samples than this dataset provides, the platelet data were used solely to evaluate the signature and not to train a model. As a preliminary step, we focused on the vascular endothelial growth factor (VEGF) signalling pathway, owing to its well-established role in promoting angiogenesis and metastasis^34^. Analysis revealed that the mean mRNA expression of VEGF pathway genes was significantly elevated in patients with metastatic cancer compared to healthy controls (p < 0.01). However, when comparing patients with primary tumours to those with brain metastases, VEGF gene expression did not differ significantly (p = NS) (**Figure 7A**), indicating that VEGF, although elevated in general metastatic disease, lacks the specificity required to distinguish brain metastases from primary tumours.

**Figure 7.**
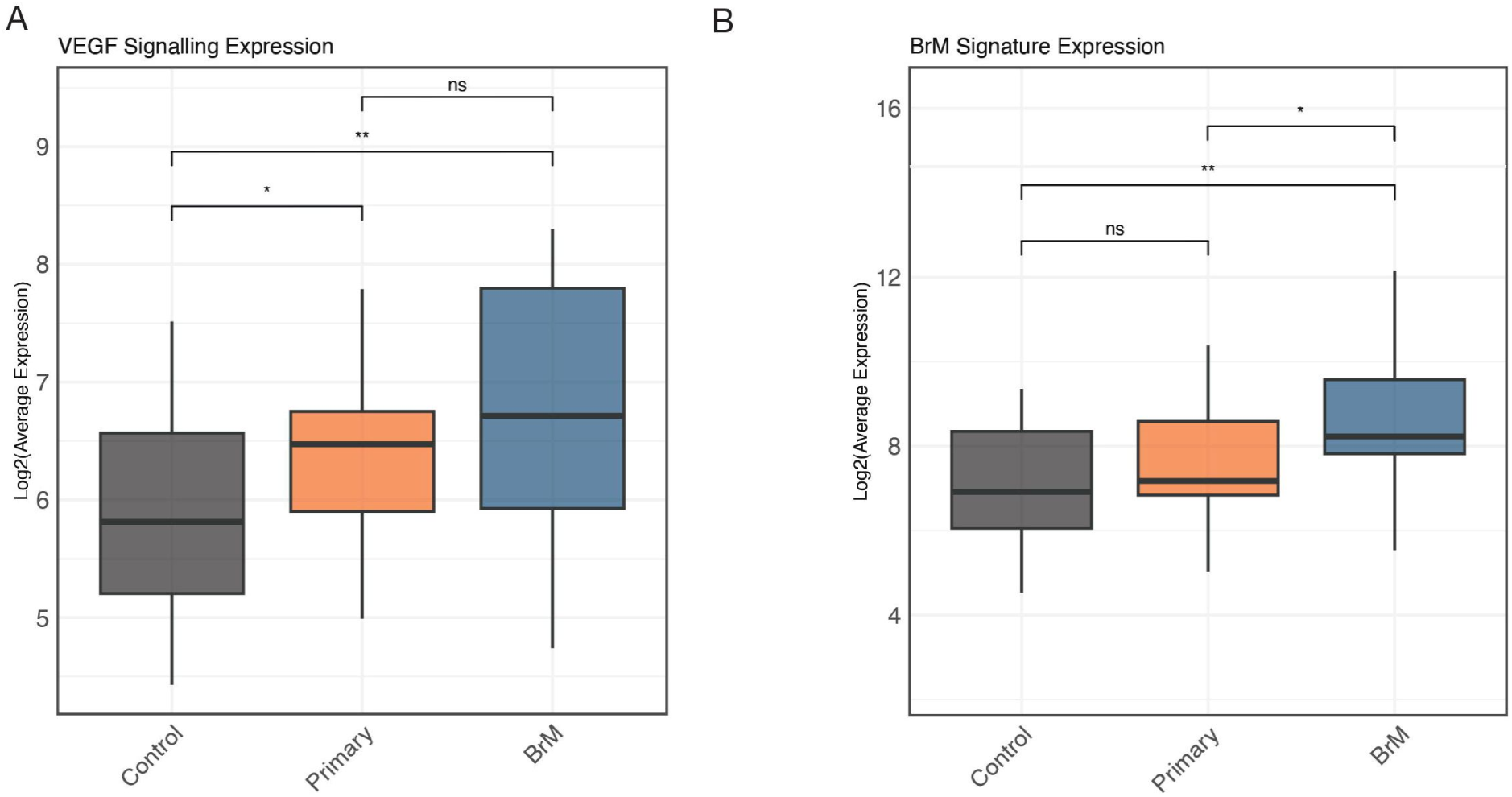
BrM signature aligns with metastatic progression trajectory and is detectable in tumour-educated platelets (TEPs). **(A)** Box plots comparing the expression levels of VEGF pathway genes in bulk RNA-seq data from TEPs of metastatic patients, primary tumour-only patients, and healthy controls across six cancer types. Significance determined by t-test (* p < 0.05, ** p < 0.01, ns = not significant). **(B)** Box plots comparing the expression levels of the 20 BrM signature genes in bulk RNA-seq data from TEPs across the same patient groups as in (D). Significance determined by t-test (* p < 0.05, ** p < 0.01, ns = not significant).

We next assessed the performance of our 20-gene BrM signature in the same dataset. Remarkably, all 20 BrM genes were significantly upregulated in TEPs from patients with brain metastases compared to both healthy donors (p < 0.01 for each gene) and patients harbouring only primary tumours (p < 0.05 for each gene) (**Figure 7B**). Importantly, none of the BrM signature genes showed differential expression between the primary tumour and healthy groups (p = NS for all), underscoring the specificity of the signature for metastatic brain involvement.

These findings demonstrate that, unlike VEGF, the BrM signature yields a robust, metastasis-specific transcriptional signal in circulating platelets. This distinct molecular profile enables accurate discrimination between brain-metastatic and non-metastatic cases, thereby positioning the BrM signature as a promising candidate for a blood-based biomarker of metastatic progression in a clinically relevant liquid biopsy context. They underscore its translational potential for early detection, prognosis, and monitoring of metastatic brain disease in cancer patients, offering a powerful, non-invasive tool for guiding therapeutic decision-making and surveillance.

## Discussion

Brain metastases (BrMs) represent a difficult clinical challenge in oncology. They are the most common intracranial tumours in adults and occur in an estimated 9-17% of all cancer patients^35^. Notably, BrMs arise most frequently from lung cancers, breast cancers, and melanomas, these three primaries collectively accounting for roughly 70% of all BrM cases^35^. The prognosis of patients with BrM is poor, and early detection is notoriously difficult because there are no routine brain metastasis screening protocols and current imaging modalities have limited sensitivity^36^. Consequently, many BrMs go undiagnosed until they become symptomatic, at which point therapeutic options are largely palliative. This clinical reality underlines the urgent need for improved strategies to predict and detect BrMs at a stage when intervention might substantially improve patient outcomes^37^.

A central insight from our study is the power of a multi-cancer analytical framework to uncover shared mechanisms underlying brain metastasis (BrM), beyond tumour-specific constraints. Prior efforts to define metastasis-associated transcriptional programs have largely been restricted to individual tumour types, limiting their generalisability across malignancies^38,39^. In contrast, recent evidence suggests that metastatic lesions arising from distinct primaries can converge on common gene expression states^40,41^. Guided by this principle, we integrated scRNA-seq data from multiple tumour types to identify a BrM-associated gene signature that transcends tissue of origin^42^. We applied ScaiVision’s supervised representation learning framework to train neural networks that classify BrM status at single-cell resolution. This approach learns tumour-agnostic low-dimensional representations that maximise separation between BrM and non-BrM cells, while preserving biological granularity. Previously reported signatures and programmes, including those described in recent pan-cancer atlases^55^, were derived for individual tumour types or with different approaches. We did not benchmark our signature against them, as such comparison lay outside the scope of this study, but it is an important next step.

Unlike conventional cluster-based or pseudobulk analyses, this method avoids averaging artefacts and enables interpretable models. Feature attribution using integrated gradients yielded a robust set of BrM-associated genes, ranked by their contribution to classification. Importantly, the resulting signature was reproducible across independent cohorts and computational strategies, supporting its generalisability. Collectively, these findings establish that a robust, cross-cancer BrM transcriptional programme can be derived from single-cell data, offering a framework for mechanistic dissection and translational development.

Scoring cells across our integrated dataset confirmed that the BrM signature is broadly conserved across tumour types and cellular compartments. Brain metastases from diverse primaries, lung, breast, renal, converged on a shared transcriptional programme, underscoring a common set of molecular adaptations required for brain colonisation. Despite distinct genetic lineages, metastatic cells consistently upregulated genes linked to angiogenesis, extracellular matrix (ECM) remodelling, and intercellular signalling. Importantly, BrM signature genes extended beyond tumour-intrinsic expression. Many encode secreted ligands or receptors mediating tumour-microenvironment communication. Ligand-receptor interaction analysis revealed robust crosstalk between malignant and host brain cells, with pro-angiogenic signalling, particularly via the VEGF pathway, emerging as a recurrent axis. This interactional footprint was conserved across metastases, suggesting a generalised mechanism of stromal engagement. Notably, VEGF family members such as VEGFA and PlGF were consistently upregulated, implicating VEGF, VEGFR1 signalling in endothelial disruption^43^ and blood-brain barrier (BBB) permeability^44,45^, consistent with prior studies. These signals are known to trigger MMP-mediated ECM degradation and junctional disassembly, facilitating tumour extravasation into brain parenchyma. Our signature’s enrichment for angiogenesis and ECM remodelling aligns closely with established mechanisms of metastasis^46^, reinforcing its biological relevance and potential utility in identifying pan-cancer vulnerabilities in BrM.

To further validate and contextualise the BrM signature, we integrated spatial transcriptomics and external datasets. Spatial profiling of brain metastasis tissue showed that the signature is expressed beyond the tumour core, including in inflammatory and vessel-associated regions, consistent with tumour-host interaction niches. VEGFA and VEGFR1 were expressed in largely distinct, neighbouring spots, as expected for paracrine signalling, providing spatial support for the ligand-receptor crosstalk inferred from single-cell data, although the spot-level resolution does not permit these signals to be assigned to specific cell types. These patterns underscore the in-situ relevance of the BrM programme and its role in shaping the metastatic niche. Cross-cohort validation using independent scRNAseq datasets from matched primary and brain metastatic samples demonstrated consistent upregulation of BrM signature genes in brain lesions across cancer types. Signature scores robustly distinguished metastases from their primary counterparts, confirming the signature’s generalisability and specificity for the metastatic state.

To explore the BrM signature’s clinical utility, we assessed its detectability in peripheral blood. Transcriptomic analysis of tumour-educated platelets (TEPs) revealed clear expression of signature genes in patients with brain metastases, but not in non-metastatic controls, supporting the notion that circulating platelets capture tumour-derived signals^47^. While VEGF-related transcripts were consistently enriched, they lacked sufficient specificity. By contrast, the broader BrM signature reliably distinguished brain metastases from primary tumours across cancer types, reflecting a distinct transcriptional programme associated with brain tropism. These observations highlight the potential of the BrM signature as a non-invasive biomarker. Its expression across diverse tumour origins and enrichment in matched metastatic samples position it as a candidate for early detection and risk stratification. In particular, the ability to detect signature transcripts in TEPs offers a feasible path toward liquid biopsy-based diagnostics. Such an assay could complement imaging by identifying metastasis prior to radiographic detection and enabling real-time monitoring of disease activity behind the blood-brain barrier. Given the frequency of brain metastases in malignancies such as lung, breast, and melanoma, a platelet-based test could have broad translational relevance in guiding surveillance and treatment response.

Among high BrM-scoring cells, MYC and ETS1 emerged as dominant transcriptional regulators, highlighting their potential role in driving brain metastasis. Both factors are established oncogenic drivers, with c-Myc coordinating programmes spanning proliferation, metabolic reprogramming, immune modulation, and angiogenesis, key processes in metastatic progression^48^. ETS1, a master regulator of invasion and neo-angiogenesis, was especially prominent due to its direct activation of matrix-remodelling enzymes and VEGF signalling pathways^49^. Notably, ETS1 activity increased in parallel with BrM scores, implicating it in the gradual acquisition of metastatic traits. Elevated ETS1 activity suggests a central role in preparing the metastatic niche, particularly through ECM remodelling and vascular co-option—mechanisms critical for colonisation of the brain. ETS1 has also been linked to the induction of L1CAM, a neural adhesion molecule that facilitates tumour cell adaptation to the brain microenvironment^50^, further reinforcing its contribution to brain-tropic behaviour.

The functional relevance of this axis is underscored by our drug repurposing analysis, which identified pazopanib, a VEGF receptor inhibitor with brain-penetrant properties, as a candidate therapeutic. Pazopanib has shown intracranial activity in clinical settings, including regression of brain metastases in patients resistant to prior therapies^51^. Similar agents, such as sunitinib, have demonstrated efficacy in lung cancer with brain metastasis^52^, supporting the premise that BrM-high cells depend on angiogenic signalling for survival and expansion. These findings point to the feasibility of anti-angiogenic strategies in curbing metastatic outgrowth within the brain.

This study has several limitations. The six tumour types analysed were those for which both primary and BrM single-cell data were publicly available, so no tumour type could be withheld to test performance in unseen tumour types, and the number of BrM samples per cancer type was small (one to seven); validation in additional tumour types will be important. The BrM score is a relative measure, and no absolute threshold applicable to new datasets or cancer types was established. The signature was derived from epithelial cells only, so its application to other cell types is exploratory, and comparisons of high- and low-scoring cells across the full dataset were not adjusted for cell type composition. The spatial data were generated at spot resolution from four patients, precluding assignment of expression and signalling to individual cell types. The platelet analysis was retrospective and was not validated prospectively or in bulk tumour cohorts, and the roles of ETS1 and VEGF signalling were inferred computationally and remain to be tested experimentally. ScaiVision was not compared with simpler baseline models, although it is based on the CellCnn algorithm, which was benchmarked in its original publication^9^. Finally, other enriched pathways (ANGPT, CDH1, CADM and IGF), candidate drugs acting through other pathways, and the performance of a smaller gene subset in platelets were not investigated and warrant further study.

In conclusion, our study delineates a transcriptional programme underpinning brain metastasis, anchored by regulators such as MYC and ETS1, and captured by a robust BrM gene signature. This work provides mechanistic insight into metastatic adaptation at the brain interface and presents actionable avenues for diagnosis and treatment. Further functional studies and prospective validation will be essential to translate these insights into clinical tools for early detection and targeted intervention, ultimately improving outcomes for patients at risk of or living with brain metastases.

## Methods

### Data Acquisition and Processing

Publicly available single-cell RNA sequencing (scRNA-seq), spatial transcriptomics (ST), and bulk RNA-sequencing datasets were utilized. scRNA-seq data, primarily generated using 10x Genomics technology, were aggregated from sources detailed in Supplementary Table 1, including paired primary lung cancer and brain metastasis (BrM) samples from GSE223503. These paired samples were used for the trajectory and regulatory network analyses and were not included in the model training or validation datasets. Within the training and validation datasets, primary tumour and BrM samples were derived from different patients. ST data were obtained from GEO accession GSE179572, and bulk RNA-seq data from GSE68086. Raw UMI count matrices and associated metadata from public scRNA-seq sources were aggregated. All data are publicly available from the cited sources. After quality control, the integrated dataset comprised 501,370 cells from 115 samples, of which 126,400 were epithelial cells used for model training and validation.

### scRNA-seq Analysis

Analyses were performed in R (v4.1.1) using Seurat (v4.0). Cells were retained if they expressed >200 genes and <20% mitochondrial reads (PercentageFeatureSet, prefix=’^MT-’) Potential doublets were identified and removed using scDblFinder (v1.6.0, default parameters) Data were normalized using log-transformation (NormalizeData, method=’LogNormalize’, scale.factor=10,000). The top 2,000 highly variable genes (HVGs) were identified (FindVariableFeatures, method=’vst’) and scaled (ScaleData). Principal component analysis (PCA) was performed on HVGs (RunPCA), and the first 30 PCs were used for downstream analyses, including K-nearest neighbour graph construction (FindNeighbors) and Louvain clustering (FindClusters, resolution=0.2). Uniform Manifold Approximation and Projection (UMAP) was used for visualization (RunUMAP, input=30 PCs)

### Cell Type Annotation

Cell types were annotated using SingleR (v1.6.1), with the Human Primary Cell Atlas reference from celldex (v1.2.0).

### ScaiVision

In the first step, ScaiVision uses data augmentation through subsampling, which increases the number of usable training samples. This data is structured into multi-cell inputs, with genes on the horizontal axis and cells on the vertical axis. Next, ScaiVision initialises filter weights randomly and computes response values for each cell-filter pair via dot products. These filters serve as connections linking gene expression values to outcome predictions (Primary or BrM), enabling automatic feature selection without the need for upfront cell clustering, ensuring the approach remains unbiased and agnostic to cell types and features. In the third step, ScaiVision aggregates cell responses for whole-sample predictions, leveraging top-k pooling, a flexible method that calculates mean values over a defined top percentage of cells, allowing sensitive detection of BrM or primary cells. Subsequently, filter values are combined in the output layer through matrix multiplication, generating probabilities for each output class.

Lastly, in network training, the network undergoes training across multiple epochs using a loss function that compares predicted and actual labels, with backpropagation and parameter updates reducing loss in each batch until a specified end condition is met, yielding a trained network with high accuracy in predicting BrM or primary. Samples were divided into training and validation cohorts, stratified by class (BrM or primary), with approximately 60% of the samples in each class assigned to training and 40% to validation. This procedure was repeated across six independent Monte Carlo cross-validation splits. ScaiVision models have previously been trained to predict sample-level disease status from scRNA-seq data using a cohort of 25 samples^54^; the present dataset was substantially larger, comprising 21 BrM and 94 primary tumour samples.

### Gene Set Analysis

Feature importance scores were calculated using Integrated Gradients within the Captum framework, defining a “BrM signature”. This signature, along with other gene sets, was scored per cell using UCell (v1.0.0), which employs a rank-based Mann-Whitney U statistic^11^. For certain analyses, cells were categorized based on their metastatic score relative to other cells within the BrM samples: ‘high-scoring’ (top 20%) and ‘low-scoring’ (bottom 20%). As scores are not directly comparable across datasets, no fixed score cutoff was applied; groups were instead defined by percentile of the BrM score within each analysis. Additionally, scores were binned into deciles (10% intervals) for visualization and analysis of score distributions.

### Trajectory and Gene Switch Analysis

Pseudotime trajectory analysis was performed on paired primary lung and BrM scRNA-seq data (GSE223503) using Monocle 2 (v2.8.0) to model metastatic progression. A CellDataSet object was created (expressionFamily=negbinomial.size()). Feature selection utilized HVGs identified in Seurat. Dimensionality was reduced using DDRTree (reduceDimension, max_components=10), and cells were ordered in pseudotime (orderCells), defining primary lung tumour cells as the root state. Branch-dependent gene expression analysis was performed using GeneSwitches (v0.1.0)^14^, identifying genes significantly associated with the high BrM branch.

### Spatial Transcriptomics Analysis

ST data generated using the Visium platform from 10X Genomics (GSE179572) were processed using Seurat. Data were normalized using SCTransform. Spatially differential expression between pre-annotated regions was assessed using FindMarkers (Wilcoxon test, Benjamini-Hochberg FDR < 0.05). The metastatic score signature was mapped onto ST spots using UCell.

### Cell-Cell Interaction Analysis

Potential ligand-receptor interactions were inferred from normalized scRNA-seq data using CellChat (v1.1.3)^18^ with the CellChatDB.human database. Cells were grouped based on cell type annotation and metastatic potential score (top 20% defined as ‘high’, bottom 20% defined as ‘low’ within BrM samples). CellChat analysis was performed using default parameters to identify significant ligand-receptor pairs and communication pathways. Interaction patterns, including the number and strength of interactions, were compared between high-scoring and low-scoring metastatic cells interacting with other cell types (e.g., stromal, immune) to identify differential communication networks potentially driving metastasis. Results were visualized using netVisual_circle and other relevant functions. Ligand-receptor interactions were additionally inferred using LIANA (v0.1.5) as an independent method.

### Gene Regulatory Network (GRN) Analysis

GRNs were inferred using CellOracle (v0.10.10)^26^ following standard documentation (https://morris-lab.github.io/CellOracle.documentation/). The workflow involved GRN construction based on TF binding motifs (JASPAR database) within putative regulatory regions (hg38), integration with normalized scRNA-seq data, and network analysis/simulation.

### Bulk RNA-seq Analysis

Bulk RNA-seq reads (GSE68086) were aligned to hg38 using STAR. Gene-level counts were quantified using featureCounts. Normalisation was performed using DESeq2 (v1.40.2)^53^ (Wald test, Benjamini-Hochberg FDR < 0.05).

### Drug Repurposing Screen

*In silico* drug repurposing was performed using ASGARD^30^ with default parameters. Differential gene expression between high- and low-scoring cells was performed using limma (v3.56.2). Normalized scRNA-seq counts from cells grouped by metastatic score (low score = control) served as input, comparing against the LINCS L1000 drug perturbation database. Drug scores were calculated, identifying candidates with significant scores across relevant comparisons.

### Statistical Analysis

Statistical analyses, including differential expression, were performed in R (v4.3.1). Comparisons between groups were made using the Wilcoxon rank-sum test. P-values from multiple tests were adjusted using the Benjamini-Hochberg method (FDR < 0.05). A p-value < 0.05 was considered significant for other tests. Differential pseudobulk gene expression analysis between high- and low-scoring cells (Figure 2E, Supplementary Figure 3D) was performed using DESeq2. Counts were aggregated by cancer type within the high- and low-scoring groups, and cancer type was included as a covariate (design: cancer type + group). Differential expression was assessed using the Wald test, with p-values adjusted by the Benjamini-Hochberg method. For Supplementary Figure 1E, BrM scores were z-scored across epithelial cells and summarised for each sample as the mean z-score. BrM and primary samples were compared within each cancer type using the Wilcoxon rank-sum test, with Benjamini-Hochberg correction across cancer types. Two BrM samples without an annotated cancer type were excluded from this comparison.

## Data availability

All the datasets used for this study are publicly available.

## Supplementary figures

**Supplementary Figure 1.**
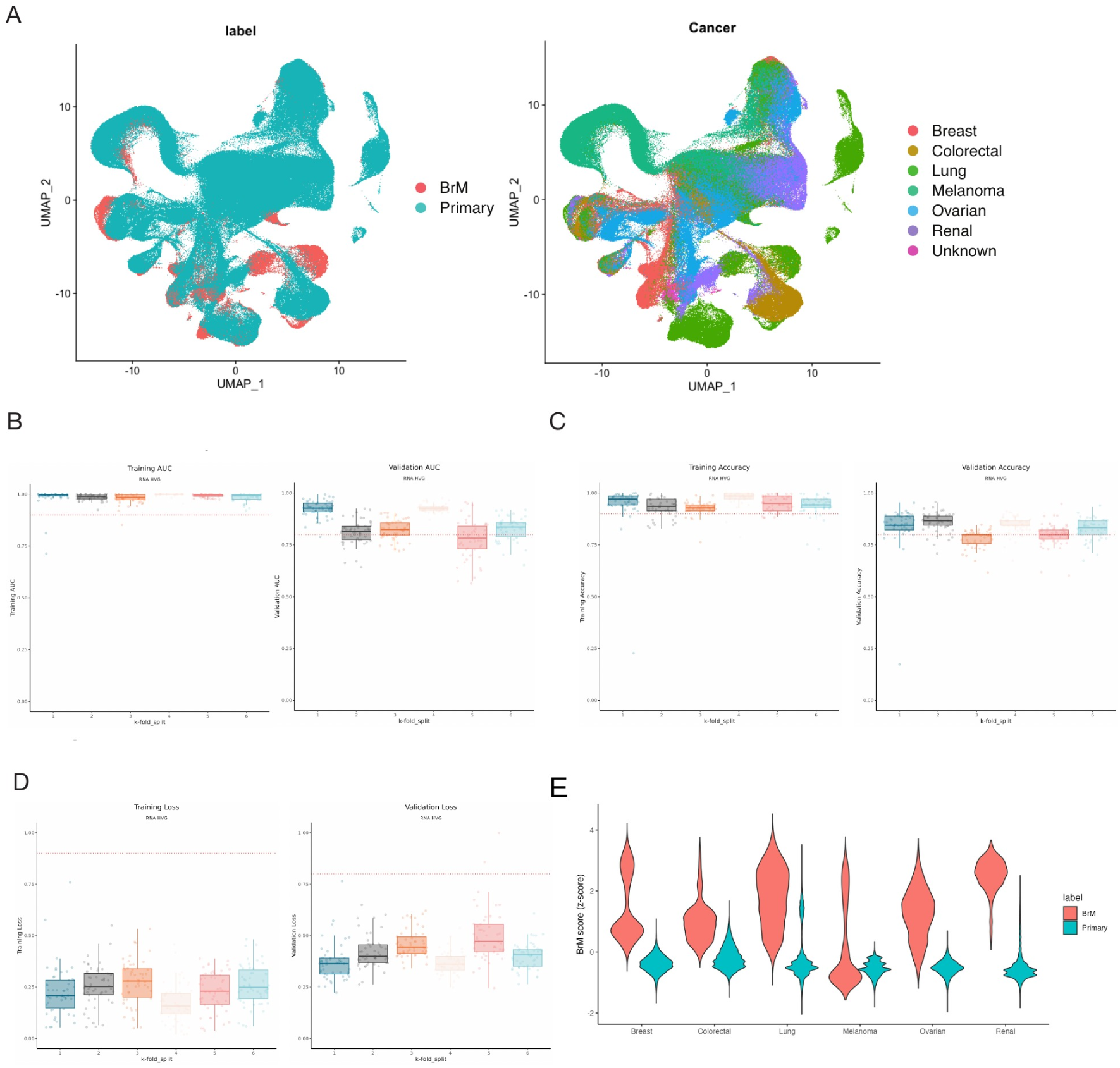
Model training and BrM signature performance in epithelial cells. **(A)** UMAP of the full dataset coloured by sample label (BrM or primary) and by cancer type. **(B)** Area under the ROC curve for training and validation samples across all folds, indicating the performance of models on training and validation data. Each point represents a separate model with its own set of hyperparameters. **(C)** Model accuracy for training and validation samples across all folds at a classification threshold of 0.5. Each point represents a separate model with its own set of hyperparameters. **(D)** Cross-entropy log-loss for training and validation samples across all folds. Each point represents a separate model with its own set of hyperparameters. **(E)** BrM signature scores (UCell, z-scored across epithelial cells) for BrM and primary samples within each cancer type. Violins represent all cells. Two BrM samples without an annotated cancer type are not shown (see Figure 2D).

**Supplementary Figure 2.**
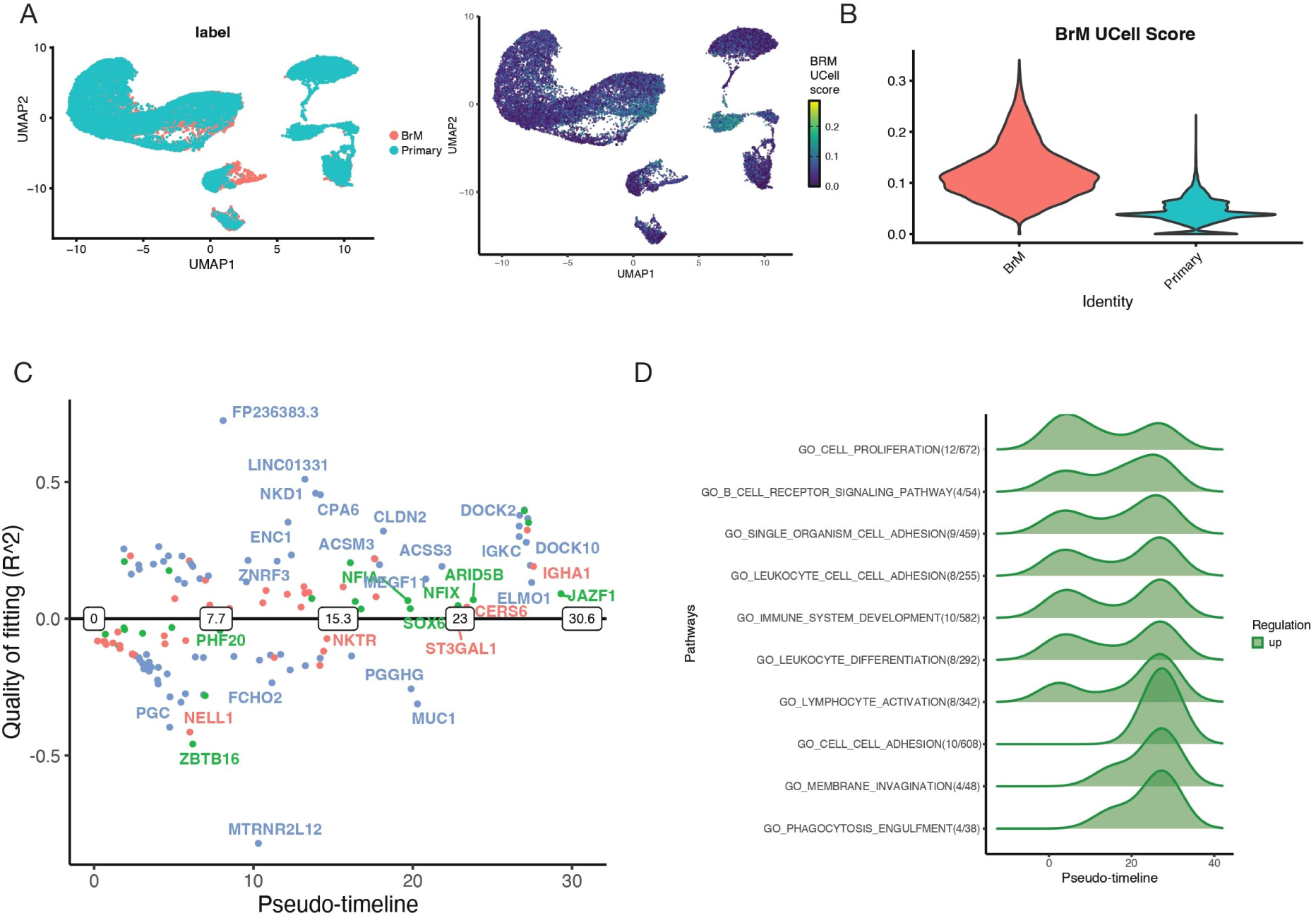
Extended BrM signature and pseudotime-based enrichment analysis. **(A)** UMAP plots of scRNA-seq data from paired primary lung and BrM samples, coloured by endpoint label, original patient ID and BrM signature score (UCell). **(B)** Violin plot highlighting higher BrM scores in BrM samples **(C)** GeneSwitches plot identifying genes significantly activated (’on’, positive R2) or inactivated (’off’, negative R2) along the pseudotime trajectory. Key genes are labelled. Colour indicates annotation of gene type. **(D)** Gene Ontology (GO) enrichment analysis results for genes dynamically regulated across pseudotime.

**Supplementary Figure 3.**
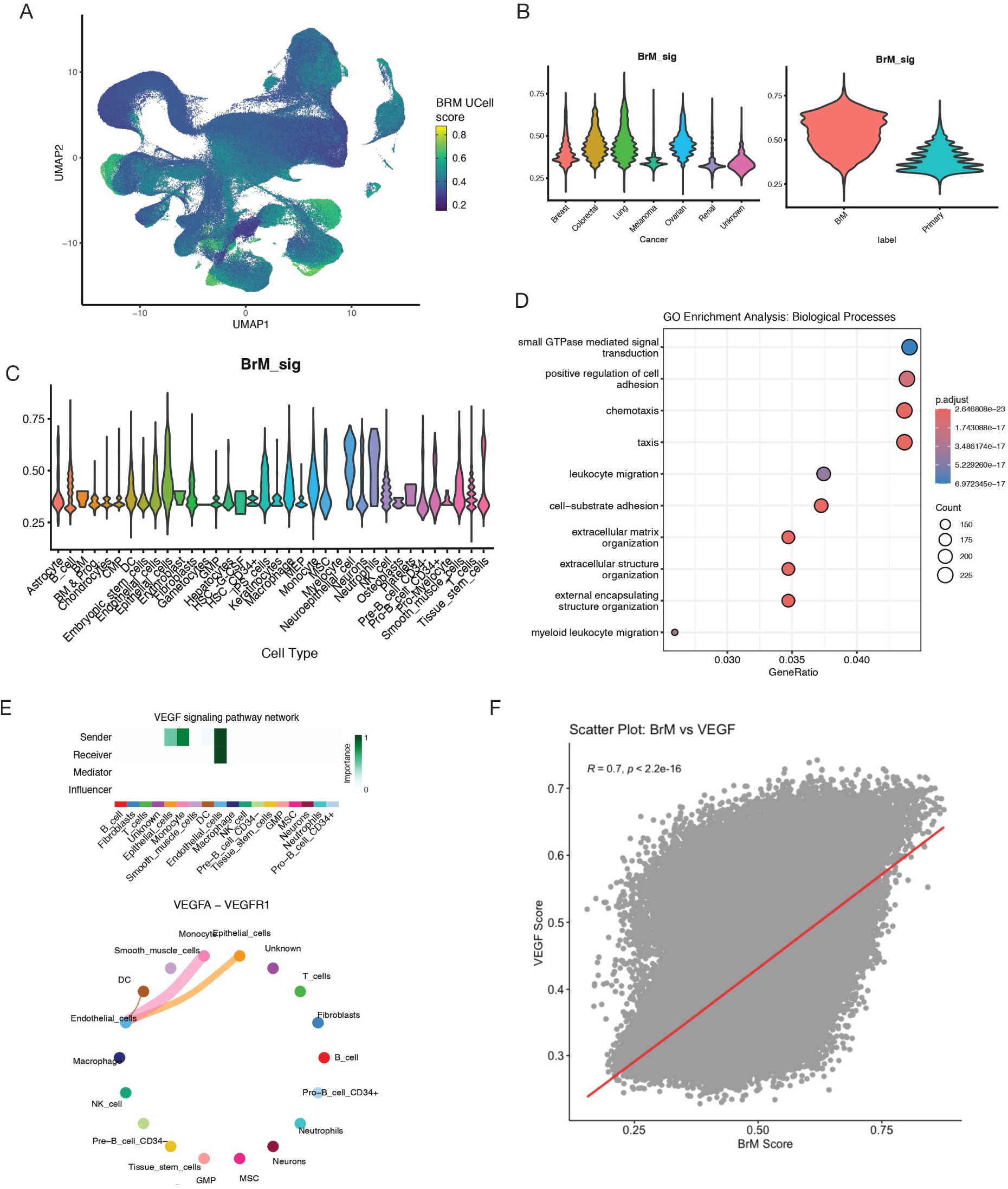
Extended cell-cell communication analysis. **(A)** UMAP plots showing BrM signature scores (UCell) projected onto all cells across multiple cancer types. Colour scale indicates BrM score. **(B)** Violin plots comparing BrM signature scores (UCell) between cancer types and endpoint labels in the multi-cancer full scRNA-seq dataset. **(C)** Violin plots comparing BrM signature scores (UCell) between cell types in the multi-cancer full scRNA-seq dataset. **(D)** Gene Ontology (GO) enrichment analysis results for biological processes upregulated in high-scoring (top 20%) compared to low-scoring (bottom 20%) cells across all cell types, based on differential pseudobulk gene expression analysis. Dot size length represents gene count; colour indicates significance level. **(E)** CellChat VEGF network visualization of the consensus cell types between low and high scored cells and ligand interactions in the consensus cell types. **(F)** Scatter plot illustrating the cell-wise correlation between BrM signature scores (UCell) and VEGF target gene set scores (UCell). Correlation coefficient (R) and p-value are indicated. Scores in this figure were computed across all cell types and are therefore not directly comparable with the epithelial-only scores in Figure 2.

**Supplementary Figure 4.**
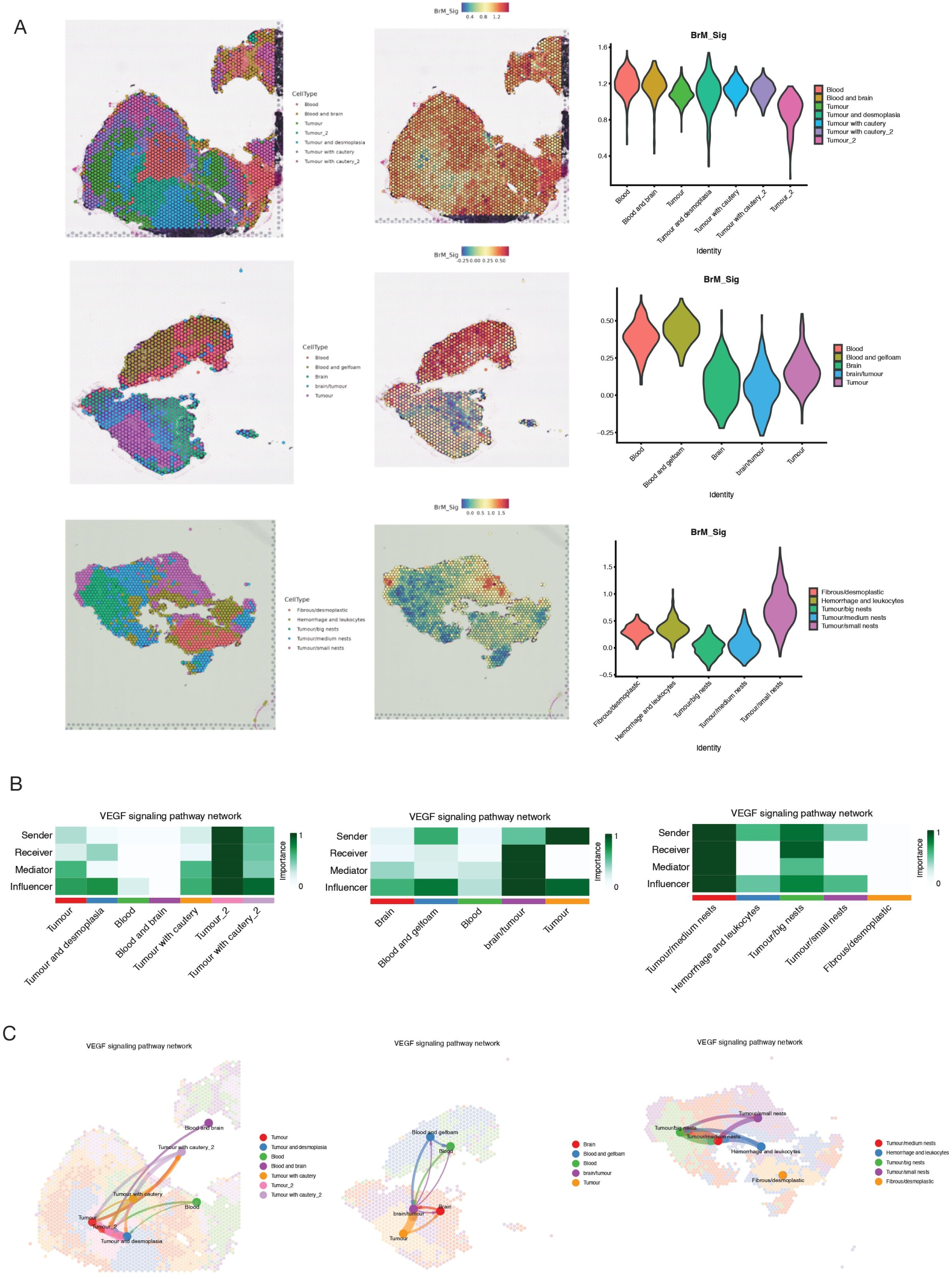
Additional spatial transcriptomics results. **(A)** Spatial feature plots showing BrM signature scores (UCell) mapped onto additional tissue sections or regions from the colorectal cancer BrM patients. **(B)** Additional CellChat network heatmap plots summarizing inferred VEGF signalling interactions between annotated tissue regions. **(C)** Additional CellChat spatial interaction plots detailing VEGF signalling interactions between annotated tissue regions in specific samples.

## Additional information

### Authors’ contributions

R. L. and V.K.T. designed the study, analyzed data, and wrote the manuscript. D.C. provided critical edits to the manuscript. S.C. co-supervised analysis and wrote the manuscript.

